# Modified methods for bovine sperm RNA isolation for consistent quality and RNA yield

**DOI:** 10.1101/2022.05.30.493985

**Authors:** Raju Kumar Dewry, Tushar Kumar Mohanty, Sapna Nath, Mukesh Bhakat, Hanuman Prasad Yadav, Rubina Kumari Baithalu

## Abstract

Sperm mRNA transcriptional profiling can be used to evaluate the fertility of breeding bulls. This study aimed to compare the modified RNA isolation methods for higher RNA yield and quality from freshly ejaculated sperm of cattle and buffalo bull for further transcriptome analysis. Ten fresh ejaculates from each Sahiwal (n = 10 bulls x 10 ejaculates) and Murrah bulls (n = 10 bulls x 10 ejaculates) were used for RNA isolation. Swim-up technique was used for live sperm separation and recovery. From the recovered live sperm, total sperm RNA was isolated by conventional methods (TRIzol, Double TRIzol), membrane-based methods combined with TRIzol (RNeasy + TRIzol) with the addition of β-mercaptoethanol (BME) and Kit (RNeasy mini) methods in fresh semen. Among different isolation methods; the membrane-based modified methods combined with TRIzol (RNeasy + TRIzol) with the addition of β-mercaptoethanol (BME) resulted significantly (P < 0.05) higher total RNA quantity (300-340 ng/μL) and better purity in different concentrations of spermatozoa viz., 30-40 million, 70-80 million and 300-400 million sperm. The study concluded that the inclusion of BME to the combined membrane-based methods with somatic cell lysis buffer solution was best for constant increased yield and purity of RNA isolation from Sahiwal cattle and Murrah buffalo bull sperm. This method will help with the interpretation of data from animal models and the consistency of clinical assessments of male factor fertility employing RNA molecular biomarkers.

## Introduction

Semen evaluation is an important criterion for successful cryopreservation and artificial insemination. Traditionally, several methods were developed to evaluate semen quality in the laboratory, like assessing genomic DNA integrity, acrosome, plasma membrane, mitochondria, and sperm-oocyte interactions. However, creating a consistent technique for routine RNA isolation from Sahiwal cattle and Murrah buffalo bull sperm to develop a non-invasive methodology to evaluate bull fertilizing ability is limited. Sperm RNA transcripts are important in spermatogenesis, sperm maturation, fertilization, oocyte genome activation, embryogenesis, and placental development, as well as the events surrounding capacitation, motility, and fertilization [1] The development of new methods such as mRNA profiling of bull sperm and understanding the significance of these transcripts would be helpful to study the fertilizing capacity of the sperm. Several investigations in humans sperm reported that spermatozoa contains more than 3,000 mRNAs that reflect the gene expression during spermatogenesis [2,3,4,5,6,7].

Earlier it was believed that spermatozoal RNAs are transcriptionally inactive; however, latest investigations reported that spermatozoa transport specific functional RNAs into the oocyte at the time of fertilization and these RNAs contribute to early embryonic development [8,9,10,11]. The process transcription is thought to be terminated after mid-spermiogenesis, and mature spermatozoa do not exhibit considerable transcriptional activity. Interestingly, distinct messenger RNA populations have been discovered in bull sperm [12] and the existence of spermatozoa RNA has been established in mammals [10,12,13,14], poultry [15] and boar [16,17]. The levels of expression of specific sperm transcripts have been linked to sperm functional characteristics [11,18,19], early embryonic development [20], and fertility [21,22]. The spermatozoa transcripts represent the spermatogenic process and have predictive value in fertilization, they may be employed as a helpful marker for high-fertile bulls [10,11,23]. To obtain precise sperm transcript expression, it is necessary to isolate sufficient quantities and quality RNA from sperm. In sperm, there is less cytoplasm, fewer full-length RNAs with physiologically degraded RNAs, and severely condensed DNA due to protamination [24]. Because of the complexity of RNA, spermatozoa require a different RNA isolation technique than other cells. To isolate total RNA, the sperm membrane and nucleo-protamine complex should be totally dissolved. Various protocols to isolate sperm RNA with varying results have been used in different species i.e., human [4,11,25,26], bovine [10,12,22,27,28], porcine [21,29] stallion [23] and chicken [15]. The lack of a suitable species-specific sperm RNA separation procedure has hindered research on sperm RNAs in the majority of species and appropriate and accurate method for sperm RNA isolation and quality assessment in Sahiwal and Murrah buffalo bull is not yet optimized. The isolation sperm RNA meets numerous challenges, specifically low RNA quantity (22-45 times less RNA than haploid spermatid) and the requirement of somatic cells removal [5,12,25]. Furthermore, sperm are extremely compacted cells with variances in sperm characteristics and chromatin packaging among species, making cellular content isolation difficult [30]. Besides, RNAs exist in the nucleus in a highly fragmented state in spermatozoa [31].

Although useful procedures for spermatozoa input were reported, an appropriate and accurate method for RNA measurement and quality assessment was not specified in detail [10,27,28]. As there are few conventional protocols for sperm RNA isolation procedures from Sahiwal cattle and Murrah buffalo bulls, this study will be key for developing a sperm RNA isolation procedure, obtaining high-quality RNA for subsequent molecular research, and presenting an effective conclusion. The purpose of this study was to develop a suitable protocol for obtaining high-quantity and high-quality RNA in indigenous cattle and buffalo and investigate a species-specific protocol for sperm RNA isolation for transcriptome analysis, regardless of seasonal variation or ejaculate quality.

## Materials and Methods

### Study location

The experiment was executed at the Artificial Breeding Research Centre, ICAR-National Dairy Research Institute (Deemed University) in Karnal, Haryana, India. The Institute is located at 29.43°N latitude and 72.2°C longitudes, at 250 meters above mean sea level. In the summer, the highest ambient temperature reaches 45°C, while in the winter, it drops to around 2°C. During July and August, the area receives 760 to 960 mm of rain, with relative humidity ranging from 41 to 85 percent.

### Experimental animal

The healthy, sexually matured Sahiwal (n=10) and Murrah buffalo (n=10) bull (4-5 years of age) which were in regular semen collection were selected for the study.

### Chemicals

The chemicals used in the experiments were procured from Sigma-Aldrich, St. Louis, MO, USA, Thermo Fisher Scientific, Waltham, Massachusetts, United States and Qiagen, Hilden, Germany.

### Semen collection and storage

Preputial washing of the bulls was carried out with normal saline one day before the semen collection. The bulls were thoroughly washed, cleaned, and dried at least 20 minutes before semen collection in the early morning. The semen sample was taken twice a week using an artificial vagina (IMV; L’Aigle, Cedex, France) set at 41°C with a 15-20 minute gap between two consecutive collections.

A total of 100 fresh ejaculates from each Sahiwal (*n*=10 × 10) and Murrah bulls (*n*=10 × 10) irrespective of seasons throughout the year. The bulls were fed with concentrate ration and seasonal green fodder as per ICAR (2013) recommendations. For quality ejaculates, followed timely vaccination, de-worming, a regular check-up for other infectious diseases, and a routine herd-health program. The fresh ejaculates were evaluated for ejaculate volume, mass motility, and initial progressive motility. Ten ejaculates were collected from each bull based on initial motility (>70%) and mass activity 3+ or higher, and one mL of neat semen was collected from each ejaculate in a 1.5-mL microcentrifuge tube for further processing.

### Sample preparation for RNA isolation

The RNA isolation area was prepared to be clean and decontaminated using RNaseZAP® spray. The following criteria were used for total RNA extraction and purification procedure [32].

✓ Free of protein (absorbance 260 nm/280 nm); undegraded (28S:18S ratio) should be roughly between 1.8 and 2.0 and accepted as pure RNA
✓ Free of genomic DNA
✓ The expected 260 nm/230 nm ratio are generally in the range of 2.0-2.2
✓ Free of nucleases for extended storage
✓ RNase free glass and plasticware
✓ The RNase and DNase free chemicals and plasticware were used for RNA isolations

### Sperm recovery and separation

#### Swim-up technique

Swim-up technique for live sperm separation and recovery was followed as per the method defined by previous researcher [33]. In brief 1 mL of freshly ejaculated semen sample was layered gently below the pre-loaded 1 mL of equilibrated sp-TALP medium in a 15 mL centrifuge tube to recover the motile spermatozoa. After preparation of the different layers (Bottom-semen sample, and Top-sp-TALP), the tubes were incubated at 39°C. After the incubation for the period of 1 hr, 750 μL of the upper fraction of the sp-TALP layer was aspirated, resuspended with 3 mL Sp-TALP medium, and centrifuged for 10 min at 300 X g. Finally, the sperm pellets were resuspended in 100 μL Sp-TALP, and the sperm recovery was estimated using a photometer (IMV, L’Aigle, France). The recovered motile sperm were varied from 300-400 × 10^6^ sperm/mL and the same concentration of sperm was used for the total RNA isolation.

#### Somatic cell removal

To ensure the purity of the sperm RNA, removal of the somatic cells must be considered during the RNA isolation procedures. A Somatic cell lysis buffer (SCLB) containing 0.1% SDS, 0.5% Triton (X-100) was used to lyse somatic cells for sperm cell purification [3,4,25,34,35].

#### Sperm concentration for total RNA isolation

The sperm concentrations of 30-40 million, 70-80 million and 300-400 million sperm cells were used for RNA isolation and comparison of different isolation methods to select the best method for achieving required quantity with quality of total sperm RNA from bull spermatozoa.

#### Isolation of total Sperm RNA

The total sperm RNA was isolated by different methods viz., conventional methods TRIzol [36], Double TRIzol, membrane-based methods combined with TRIzol (TRIzol + RNeasy) with the addition of β-mercaptoethanol (BME) [37] and pure kit methods with slight modifications.

### Conventional methods

#### TRIzol method

The pre-prepared sperm pellet was mixed with 1 mL of PBS, and washed three times by centrifugation at 3000 rpm for 5 min at 4°C for purification of the sperm cell from other somatic cells, germ cells and leukocytes. According to the manufacturer’s instructions, total sperm RNA was extracted from bull sperm using TRIzol (Invitrogen, USA) with minor modifications.

In a short, 0.5 mL of ice-cold TRI reagent was added to the sperm pellet and homogenised 25-30 times with a 20-G needle attached to a 5-mL syringe. After vortexing for 5 minutes, the samples were incubated at room temperature (RT) for 5 minutes until the sperm membrane was completely dissociated. 200 μl chloroform was added to the lysate and thoroughly shaken by hand for at least 20 seconds, followed by 5 minutes at room temperature. The mixture was centrifuged at 12,000 X g for 20 min at 4°C to separate the phases. After centrifugation, three layers were observed viz. the upper aqueous layer (RNA), middle white layer (DNA) and bottom pink layer (protein). The RNA-containing upper aqueous layer was transferred to a new 1.5 mL conical microcentrifuge tube, where an equal volume of isopropanol was added and gently mixed by reversing the tubes. The mixture was kept for 10 min at RT and centrifuged at 12,000g for 15 min at 4 °C. Supernatant was discarded after centrifugation and 1 mL of 99.99% ethanol was added to the RNA pellet and again centrifuged at 12,000g for 10 min. Finally, ethanol was removed, and the RNA pellet was air-dried. The pellet was dissolved in 40 μl of nuclease-free water, followed by the addition of 2 μl of RNase inhibitor (20 U/l, Invitrogen) and 10 mM dithiothreitol (DTT).

#### Double TRIzol method

The prepared pellet was resuspended in 1 mL of PBS, and washed three times by centrifugation at 3000 rpm for 5 min at 4°C to purify the spermatozoa by removing the other somatic cells, germ cells and leukocytes.

TRIzol reagent was used to extract total RNA from spermatozoa. In this procedure the samples were lysed twice with TRIzol reagent for complete dissociation of sperm membrane. In short, 0.5 mL of ice-cold TRIzol was added to sperm pellet and homogenised for a minimum of 25 to 30 times with a 20-G needle attached to a 5-mL syringe. The samples were then vortexed for 5 minutes at room temperature and incubated for 5 minutes, centrifuged at 12,000g for 30 seconds, 0.5 mL of fresh, chilled TRIzol was added to the supernatant and vortexed for 1-2 minutes, and the rest of the procedure was carried out as described in the TRIzol method.

### Combined method

#### RNeasy + TRIzol method with β-mercaptoethanol (BME)

The pre-prepared pellet was mixed with 1 mL of PBS, and the sperm sample was washed three times by centrifugation at 3000 rpm for 5 min at 4°C to remove the somatic and other germ cells.

Here, the total sperm RNA was isolated by combining the conventional and kit (RNeasy) methods described by earlier worker [25] with modifications (Fig. 1). Briefly, the sperm pellet was homogenised with a 20-G needle for about 25-30 times in 1 mL SCLB containing - β mercaptoethanol (10 μl/mL). The homogenized mixture was vortexed for 1-2 min and incubated at RT for 30 minutes. 0.5 mL of ice-chilled TRI reagent was added to the mixture and again subjected to vortexing for 2 minutes. The mixed sample was then maintained at room temperature for 15 minutes without being disturbed before adding 200 μl cooled chloroform. The contents were mixed vigorously by hand for 20 s and kept at RT for 3 to 5 minutes, followed by centrifugation at 12,000*g* for 20 minutes at 4 °C. The upper aqueous layer was aspirated into a sterilized 2.0-mL tube, and an equal volume of 100 percent ethanol was pipetted in and mixed thoroughly. The mixture was transferred to a mini-spin column provided and processed as per the manufacturer’s protocol. Three times elution was done at room temperature with different volumes (30, 25 and 15 μl) of nuclease-free water. Finally, the three elutes were pooled, and 10 mM DTT was added to the pooled RNA, which was stored at −80°C until further processing.

**Fig. 1.**
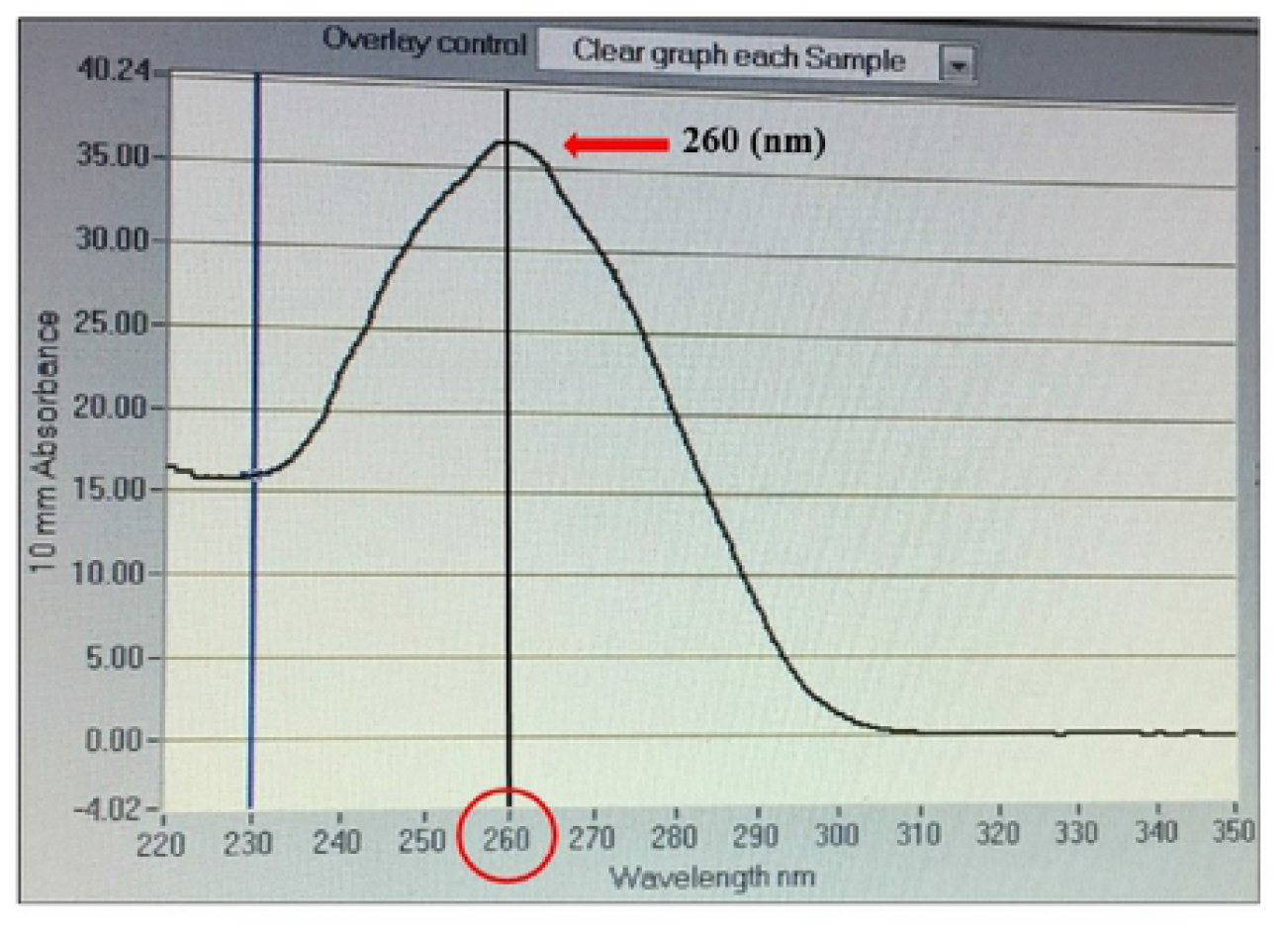
RNA isolated by the modified methods showed peak absorbance at 260 nm even with effective concentration of RNA, indicating good quality RNA

#### RNeasy + TRIzol method without β-mercaptoethanol

The pre-prepared sperm pellet was resuspended in 1 mL of PBS, and the sperm sample in PBS was centrifuged three times for 5 minutes at 3000 rpm at 4°C to separate the spermatozoa from somatic cells, germ cells, and leukocytes, as described above.

Total sperm RNA was isolated using a modified combination of conventional and RNeasy methods. BME was not used in this method, unlike the previous one [25].

### Membrane-based method

#### RNeasy mini kit method

The pre-prepared sperm pellet was resuspended in 1 mL of PBS and washed three times by centrifugation at 3000 rpm for 5 min at 4°C to remove other somatic cells, germ cells, and leukocytes from the spermatozoa.

The QIAGEN RNeasy® Mini Kit (Cat No./ID: 74104) was used to isolate the total RNA as per the manufacturer’s instructions. To eliminate genomic DNA, RNAse-free, DNAse I treatment (Qiagen) was performed.

### Estimation of total RNA yield and quality

#### Importance of RNA integrity

RNA is a very unstable molecule, which is rapidly digested by ubiquitous RNase enzyme presence. Because of the highly degradable, shorter RNA fragments frequently occur in a sample, potentially compromising downstream applications [38,39].

#### Nano Drop spectrophotometer

Estimated total RNA concentration and the optical density ratios (OD-260/280 and OD-260/230) with nono drop spectrophotometer (ND-1000, Thermo Scientific, USA). The absorbance at 230, 260 and 280 nm was used to determine the purity of isolated RNA. The 260/280 ratio was used to determine potential protein contamination, whereas the 260/230 ratio was used to determine salt and organic solvent contamination. The absorbance at 260 nm was measured with 1μL of RNA sample, and the total RNA concentration was represented in nanograms per microliter (ng/μl) units. A bioanalyzer was used to examine the RNA fragment size distribution and peak (Agilent 2000, Agilent Technologies, USA).

#### Gel Electrophoresis

The quality of the isolated RNA was observed by gel electrophoresis using 2.0% agarose gel stained with ethidium bromide to visualize the double band rRNA. Electrophoretic methods have been applied to separate the samples according to the size of comprised molecules to evaluate the degree of degradation. Reliable RNA integrity is evaluated using agarose gel electrophoresis stained with ethidium bromide, which produces a specific band pattern [40]. Gel image shows double bands containing the 28S and 18S ribosomal RNA (rRNA) and other bands. When the ratio of 28S:18S bands is within 1.8 to 2.0 at 260/280 nm, the RNA sample is considered as high quality.

#### Statistical analysis

The data recorded were analyzed using the SPSS software (version 22). One-way analysis of variance (ANOVA) followed by Tukey’s post hoc test to compare the means and determine the significant differences between the groups were used to select the input spermatozoa concentration and the protocol used for RNA isolation based on total RNA yield and quality. The values were presented as means ± SE, and p < 0.05 was considered to be statistically significant.

## RESULTS

### Sperm purification and recovery

A swim-up technique was applied to the freshly ejaculated semen sample to recover the motile spermatozoa. The recovered motile sperm was 300-400 × 10^6^ sperm/mL and different concentrations of sperm *viz*., 30-40, 70-80 and 300-400 million sperm cells were used for the RNA isolation. The results of total RNA yield and the purity of isolated RNA using different protocols are described below-

### Total Sperm RNA yield

According to spectrophotometric measurement, total RNA yield in Sahiwal and Murrah buffalo bull spermatozoa was considerably (p<0.05) higher in the TRIzol + RNeasy procedure with the addition of BME in 300-400 million sperm concentrations than in all other methods under consideration (Table 1 and 2).

**Table 1.** Total RNA yield of Sahiwal bull spermatozoa using different isolation methods (*N*=100) (Mean ± SE)

| S.No. | RNA Isolation Method | Total RNA yield (ng/μL) | | |
| --- | --- | --- | --- | --- |
| | | Sperm Input Concentration (300-400 x $10^6$) | Sperm Input Concentration (30-40 x $10^6$) | Sperm Input Concentration (70-80 x $10^6$) |
| 1 | TRIzol | 224.70 ± 10.28<sup>b</sup> | 15.76 ± 0.36<sup>bc</sup> | 21.38 ± 0.41<sup>b</sup> |
| 2 | Double TRIzol | 238.48 ± 11.32<sup>abc</sup> | 24.49 ± 0.52<sup>bd</sup> | 30.31 ± 0.55<sup>b</sup> |
| 3 | RNeasy | 52.29 ± 4.24<sup>c</sup> | 8.88 ± 0.22<sup>c</sup> | 12.01 ± 0.27<sup>cd</sup> |
| 4 | TRIzol + RNeasy | 304.51 ± 14.61<sup>abc</sup> | 29.87 ± 0.37<sup>b</sup> | 45.89 ± 0.74<sup>bcd</sup> |
| 5 | TRIzol + BME + RNeasy | 340.72 ± 19.31<sup>a</sup> | 39.24 ± 0.41<sup>ab</sup> | 68.87 ± 1.75<sup>abc</sup> |

Means bearing superscripts (a,b,c,d) within a column for a particular parameter differ significantly ( $P < 0.05$ )

**Table 2.** Total RNA yield of Murrah bull spermatozoa using different isolation methods (*N*=100) (Mean ± SE)

| S.No. | RNA Isolation Method | Total RNA yield (ng/μL) | | |
| --- | --- | --- | --- | --- |
| | | Sperm Input Concentration (300-400 x $10^6$) | Sperm Input Concentration (30-40 x $10^6$) | Sperm Input Concentration (70-80 x $10^6$) |
| 1 | TRIzol | 207.46 ± 3.83<sup>ab</sup> | 19.74 ± 0.57<sup>c</sup> | 23.07 ± 0.59<sup>bd</sup> |
| 2 | Double TRIzol | 222.02 ± 5.75<sup>ab</sup> | 27.86 ± 3.14<sup>b</sup> | 32.68 ± 0.63<sup>b</sup> |
| 3 | RNeasy | 46.06 ± 2.44<sup>cd</sup> | 12.11 ± 0.47<sup>bd</sup> | 15.35 ± 0.43<sup>c</sup> |
| 4 | TRIzol + RNeasy | 261.38 ± 4.83<sup>ac</sup> | 37.39 ± 1.04<sup>abc</sup> | 41.73 ± 0.70<sup>bcd</sup> |
| 5 | TRIzol + BME + RNeasy | 318.20 ± 37.60<sup>a</sup> | 51.12 ± 2.16<sup>ac</sup> | 72.23 ± 2.65<sup>abc</sup> |

Means bearing superscripts (a,b,c,d) within a column for a particular parameter differ significantly ( $P < 0.05$ )

The TRIzol + RNeasy + BME method resulted in the highest total RNA yield of 39.24 ± 0.41ng/μL, 68.87 ± 1.75ng/μL and 340.72 ± 19.31, ng/μL in 30-40 × 10^6^, 70-80 × 10^6^ and 300-400 × 10^6^ sperm input concentration, respectively as compared to other isolation methods in Sahiwal bull sperm. It is followed by the TRIzol + RNeasy method, double TRIzol method, TRIzol, and RNeasy (Table 1). Similar results were found in the Murrah bull sperm with the highest total RNA yield in TRIzol + RNeasy + BME method resulting 51.12 ± 2.16 ng/μL, 72.23 ± 2.65 ng/μL and 318.20 ± 37.60 ng/μL in 30-40 × 10^6^, 70-80 × 10^6^ and 300-400 × 10^6^ sperm input concentration, respectively followed by TRIzol + RNeasy method, double TRIzol method, TRIzol and RNeasy (Table 2).

### Sperm RNA quality

The TRIzol + RNeasy with the addition of the BME method yielded good quality RNA without contaminating substances than the other methods in both Sahiwal and Murrah bull spermatozoa (Table 3 and 4), the peak at 260 mm absorbance (Fig. 1). In Sahiwal bull sperm, the isolated RNA’s purity was best in the TRIzol + BME + RNeasy method with 1.95 ± 0.00 ng/μL, 1.93 ± 0.01 ng/μL and 1.90 ± 0.01 ng/μL in 30-40 × 10^6^, 70-80 × 10^6^ and 300-400 × 10^6^ sperm input concentration, respectively followed by TRIzol + RNeasy, RNeasy, TRIzol and Double TRIzol (Table 3). Similar trends also resulted in Murrah bull sperm. The best quality RNA was found in TRIzol + BME + RNeasy method with 1.97 ± 0.00 ng/μL, 1.91 ± 0.01 ng/μL and 1.98 ± 0.00 ng/μL in 30-40 × 10^6^, 70-80 × 10^6^ and 300-400 × 10^6^, respectively followed by TRIzol + RNeasy, RNeasy, TRIzol and Double TRIzol (Table 4).

**Table 3.** RNA purity of Sahiwal bull spermatozoa using different isolation methods (*N*=100) (Mean ± SE)

| S.No. | RNA Isolation Method | RNA purity<br>Absorbance at 260/280 |  |  |
| --- | --- | --- | --- | --- |
|  |  | Sperm Input Concentration<br>(300-400 x 10<sup>6</sup>) | Sperm Input Concentration<br>(30-40 x 10<sup>6</sup>) | Sperm Input Concentration<br>(70-80 x 10<sup>6</sup>) |
| 1 | TRIzol | 1.67 ± 0.01<sup>d</sup> | 1.65 ± 0.01<sup>cd</sup> | 1.68 ± 0.01<sup>d</sup> |
| 2 | Double TRIzol | 1.66 ± 0.02<sup>c</sup> | 1.77 ± 0.01<sup>d</sup> | 1.77 ± 0.01<sup>cd</sup> |
| 3 | RNeasy | 1.83 ± 0.01<sup>ad</sup> | 1.9 ± 0.00<sup>abc</sup> | 1.85 ± 0.01<sup>ac</sup> |
| 4 | TRIzol + RNeasy | 1.87 ± 0.01<sup>ab</sup> | 1.88 ± 0.00<sup>ac</sup> | 1.83 ± 0.00<sup>ab</sup> |
| 5 | TRIzol + BME + RNeasy | 1.90 ± 0.01<sup>ac</sup> | 1.95 ± 0.00<sup>ab</sup> | 1.93 ± 0.01<sup>a</sup> |

Means bearing superscripts (a,b,c,d) within a column for a particular parameter differ significantly ( $P < 0.05$ )

**Table 4.**
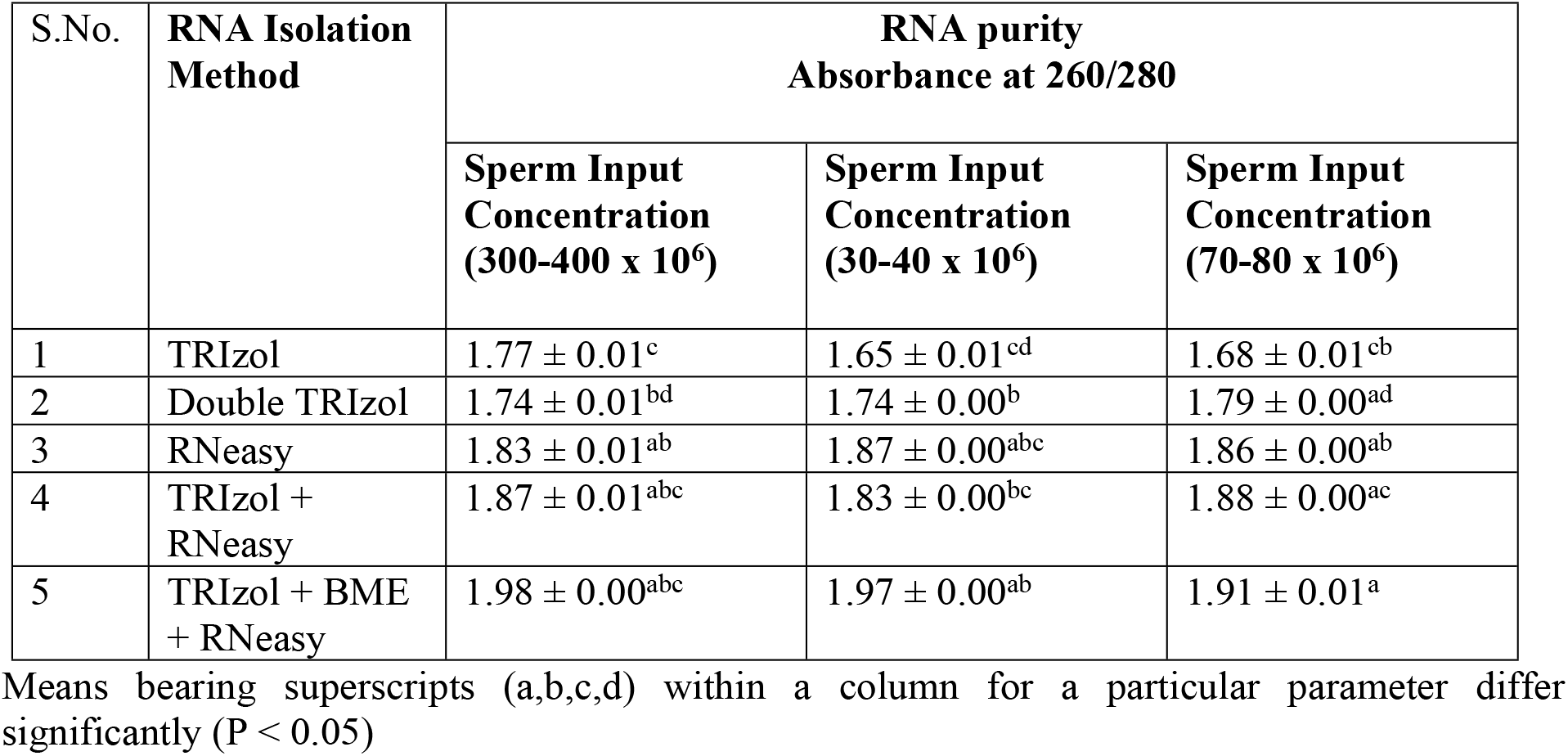
RNA purity of Murrah bull spermatozoa using different isolation methods (*N*=100) (Mean ± SE)

The Agilent 2100 Bio-Analyzer was used for RNA integrity number (RIN) to estimate the purity or integrity of RNA. Based on the peaks, the RNA integrity was measured by RNA integrity number value. RIN values are measured from 1 - 10, where RIN Value 1 - 5 indicates complete degradation and 5 - 7 shows partially degraded RNA and RIN value above 7 indicates high quality RNA. In our study, the RIN value of 2.1 to 9.7 was observed in different samples, indicating the RIN of 9.7, highly intact and 2.1 as highly degraded (Fig.2). Persistence of 18S and/or 28S rRNA showing the isolated RNA integrity (Fig.2). The double band rRNA was visualized by running 2.0% agarose gel electrophoresis (Fig. 3).

**Fig. 2.**
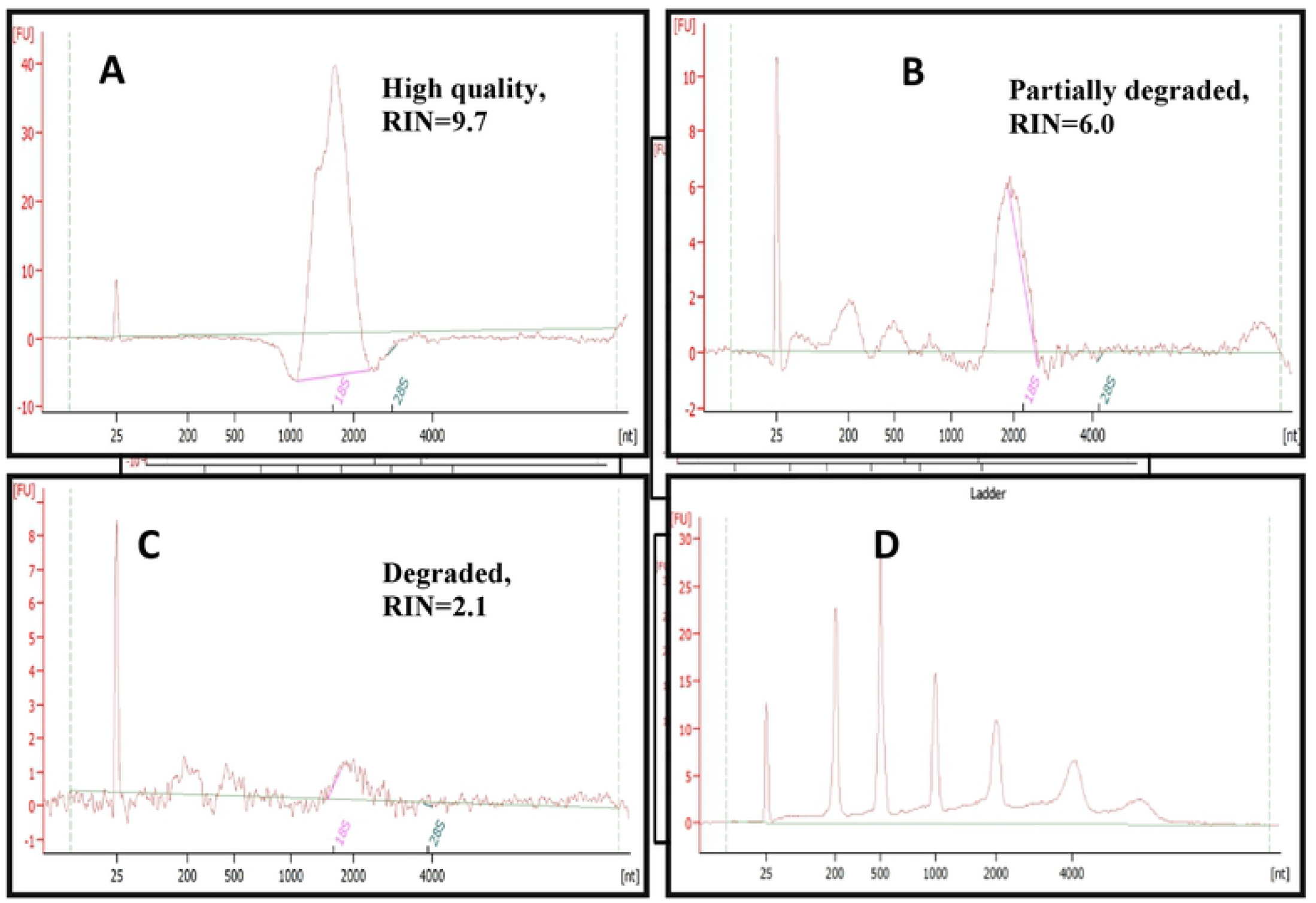
RNA Integrity Value at different grades: (A) highly intact RNA, (B) partially <u>partially</u> degraded, (<u>C</u>C) D<u>d</u>egraded RNA and (D) the nucleotide size was calculated using the s<u>S</u>tandard marker.

**Fig. 3.**
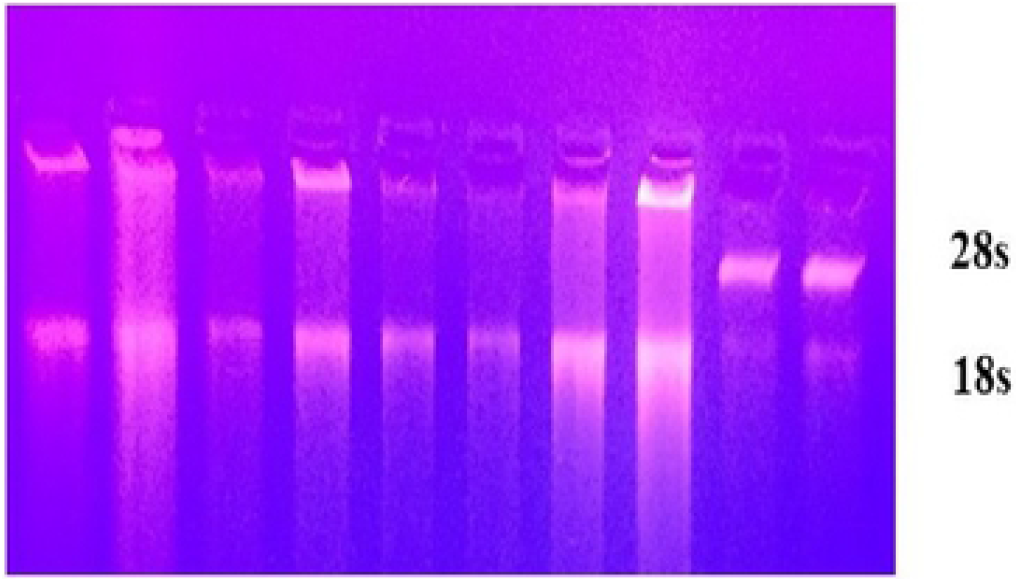
Double band rRNA was visualized by 2.0% agarose gel electrophoresis

## Discussion

In the present study, we considered the motile and spermatozoa free from other contaminating and somatic cells. The total RNA yield was highest in the modified method (TRIzol + RNeasy + BME), i.e., 39.24 ± 0.41ng/μL, 68.87 ± 1.75ng/μL and 340.72 ± 19.31, ng/μL in 30-40 × 10^6^, 70-80 × 10^6^ and 300-400 × 10^6^ sperm input concentration, respectively as compared to other isolation methods in Sahiwal bull sperm. It was followed by the TRIzol + RNeasy method, double TRIzol method, TRIzol, and RNeasy (Table 1). Similar trends were found in the Murrah bull sperm with the highest total RNA yield in TRIzol + RNeasy + BME method resulting 51.12 ± 2.16 ng/μL, 72.23 ± 2.65 ng/μL and 318.20 ± 37.60 ng/μL in 30-40 × 10^6^, 70-80 × 10^6^ and 300-400 × 10^6^ sperm input concentration, respectively followed by TRIzol + RNeasy method, double TRIzol method, TRIzol and RNeasy. The same method displayed the best optical density (OD) at A260/A280 ratio of 1.97, 1.93 and 1.90 in 30-40 × 10^6^, 70-80 × 10^6^ and 300-400 × 10^6^ sperm input concentration, respectively in Sahiwal bull sperm, and 1.97, 1.91 and 1.98 in 30-40 × 10^6^, 70-80 × 10^6^ and 300-400 × 10^6^ sperm input concentration, respectively in Murrah bull sperm, which indicates that samples free from protein and organic-free substances among the different methods used. In earlier studies [3], mammalian spermatozoa gene expression profiling has been projected as a novel non-invasive tool to evaluate male fertility. Regardless of the acute contribution of sperm RNA toward the male fertility [5,41], embryo development [25], epigenetic inheritance for acquired traits involved in the paternal genome [42] and health [43], the scarcity of optimization of transcriptomic investigation outfits prevents a thorough understanding of sperm biology. This research aimed to recommend a methodology for extracting high-quality total RNA from bull spermatozoa for transcriptome analysis. Because of reduced cytoplasm, a low number of intact and physiologically degraded RNAs [11], a high proportion of gDNA, and protamination of the nucleus in sperm, optimising an RNA separation process for bovine sperm is crucial [25]. Because the accuracy of functional genomics research is determined by RNA quality and inter-species variability in sperm characteristics and packaging, sperm RNA separation procedures must be adapted to each species.

Even after the purification of spermatozoa by suitable sperm separation techniques, a major challenge is to extract RNA of sufficient quantity and high quality RNA (i.e., non-degraded RNA and decontaminated). Sperm needs to be purified using various methods for making sperm RNA free from other cell contaminants, such as swim-up protocol [29,44], percoll gradient solution [12,28] and somatic cell lysis [3,11,26]. The different procedures of sperm separation and semen storage condition might influence the sperm recovery and the total RNA yield [24,45].

Throughout the sperm epididymal maturation, sperm protamines undergo thiol oxidation intra-molecular form followed by inter cell disulfide bonds [46]. The covalent sulfur-sulfur (S-S) bonds stabilize the sperm DNA and are believed critical to condense the mammalian sperm nucleus into its fully mature form. During the period of RNA isolation, utmost care should be adopted to avoid degradation by RNases enzymes. Intracellular RNases are released during the RNA isolation procedure, particularly at the lysis step and must be rapidly and thoroughly inactivated to obtain high-quality RNA. β-mercaptoethanol is deliberated a reducing agent that conclusively denatures RNases by disrupting the disulfide bonds and by removing the innate conformation required for enzyme action. Any RNases present in the sample to be extracted will be completely deactivated when combined with the strong denaturing properties of guanidiumisothiocyanate (GITC) and RLT buffer supplied in the RNeasy kits [47]. The addition of β-mercaptoethanol can act as a biological antioxidant by scavenging hydroxyl radicals and might be resulted in a high yield of RNA because of its property to break the disulfide bonds (S-S) and additional lysis and nuclear component dissociation. Since membranes provided in the kit can hold only RNA, repeated washing procedures eliminate other contamination like proteins, salts, and inorganic solvents.

As a major source of bacteria found in soil, bedding, and manure, the bull’s preputial orifice is one of the most important sources [48] and to minimize the bacterial load from the ejaculates before collection of the semen; preputial washing was done with normal saline. Preputial wash helps yield good quality RNA since the ejaculates collected for RNA extraction were free from any contamination like somatic cells, etc. Somatic cell lysis buffer is one of the potent buffer which helps in the lysis of any somatic cells. The composition of somatic cell lysis buffer and the time the sperm cells are exposed to lysis buffer are also very important while isolating spermatozoal RNA, since more prolonged exposure removes the somatic cells, yielding high quality RNA profiles in sperm samples. RNA quantity and quality were found to be far better in the case of TRIzol associated with membrane-based methods (TRIzol + RNeasy), supported by earlier reports [37,49] when compared to membrane-based RNA isolation methods alone. The TRIzol reagent protects RNA integrity while breaking cells and dissolving cell components during the homogenization or lysis process. The quality of RNA was better in the membrane-based kit method resulted in our study also in agreement with previous reports [15].

Even though the overall structure of spermatozoa is similar across species, very combative procedures must be required to disrupt the membrane structure of bovine spermatozoa. By heating the lysis buffer supplied in the Qiagen kit, human sperm RNA samples were able to extract [3,4]. This method was ineffective for bull spermatozoa because extensive cellular debris is deposited at the bottom of the tube after the lysis process, resulting in low total RNA quantity. Indeed, mature spermatozoa are not known to be transnationally active, implying that rRNAs required for ribosome assembly may not be present.

Five techniques have been attempted and compared to recommend a suitable method for bovine sperm RNA isolation. When membrane-based approaches combined with TRIzol (TRIzol + BME + RNeasy) were compared to other methods, both the amount and quality of recovered RNA were higher (TRIzol, Double TRIzol and RNeasy). Our study’s findings agree with earlier researchers [37] who isolated RNA from Holstein Friesian bulls and reported higher RNA yield in RNeasy + TRIzol method compared with the other methods. Higher concentration (37.39 ± 1.04 ng/μL) was found in the present study from 30-40 million sperm as compared to the previous research (15.22 ± 0.27 ng/μL in 30 million sperm) in Murrah Buffalo bull sperm using heated TRIzol + RNeasy mini method with PVP-coated silica colloidal solution (PVA-Si; Percoll) [49]. Similarly another study reported using TRIzol + RNeasy method with an average RNA yield of 8.15 ng/μL from 70-80 million sperm in Gir bull, which was found to be lower than the present investigation 68.87 ± 1.75 ng/μL and 72.23 ± 2.65 in Sahiwal and Murrah bull sperm, respectively [50]. This might be due to μ -mercaptoethanol effectively dissolved the nucleoprotamine complex and unfolded the nuclear proteins, resulting in improved nuclear component lysis and dissociation and the elimination of gDNA in the combined lysis procedures. Additionally, the purity results found in the kit method, which is membrane-based, specifically holds the only RNA; hence, the RNA’s purity was superior with no/less gDNA in the membrane-based techniques. Moreover, the repeated washing procedure removed the salts and organic solvents in the membrane-based methods and demonstrated a peak at 260 nm using a spectrophotometer. The RIN value and the peaks observed in the present study with the value of 2.1 to 9.7 was pragmatic in different samples and are also following the earlier reports [49,50,51].

In conclusion, our findings suggest that the membrane-based methods with TRIzol and somatic cell lysis buffer, BME or DTT, and for generating good quality and quantity RNA from fresh bovine and buffalo bull spermatozoa, further phase separation is necessary and highly effective. We have observed consistent RNA yield and quality from the 100 ejaculates from both cattle and buffalo bull in our findings irrespective of seasonal variation of semen quality which may help further transcriptome analysis and other molecular studies of semen biology. This is the first report of consistent of RNA yield and quality from fresh ejaculated semen from Sahiwal cattle and Murrah bull irrespective of seasonal variation of semen quality. However, further optimization of RNA yield and quality from frozen semen is required from indigenous cattle and buffalo.

## ACKNOWLEDGMENTS

The authors sincerely acknowledge the Director cum Vice-Chancellor, ICAR-National Dairy Research Institute, Karnal, Haryana, India, to provide the necessary facilities to carry out this work. The authors are thankful to Dr. A. K. Mohanty, Principal Scientist and Dr. T. K. Datta, Principal Scientist, Animal Biotechnology Centre, ICAR-National Dairy Research Institute for providing NanoQuant and NanoDrop spectrophotometer for quantification of the RNA yield and thankful to Dr. S. De, Principal Scientist and Head, Animal Biotechnology Centre for his valuable suggestion and guidance for the work.

## ETHICAL APPROVAL

The Institutional Animal Ethics Committee approved all experimental procedures (43-IAEC-18-21).

## Author Contributions

**R.K. Dewry**: Methodology, Writing-Original draft preparation, Formal analysis. **T.K. Mohanty:** Conceptualization, Resources, Funding acquisition, Methodology, Supervision, Investigation, Reviewing and Editing, Final approval of the version to be submitted. **S.Nath**: Methodology, Investigation, Formal analysis, Reviewing and Editing. **M. Bhakat**: Investigation, Supervision, Reviewing and Editing. **H.P. Yadav**: Sample collection, Reviewing and Editing. **R. K. Baithalu**: Supervision.

